# An animal model of coronary microvascular dysfunction (CMD) in the female spontaneously hypertensive rat: the role of diet

**DOI:** 10.1101/2025.01.14.632950

**Authors:** Rami S. Najjar, Puja K. Mehta, Andrew T. Gewirtz

## Abstract

Coronary microvascular dysfunction (CMD) drives angina in patients with ischemia non-obstructive coronary artery disease, a condition that is more prevalent in females. Effective treatment strategies and a detailed mechanistic understanding of CMD remain limited, in part, due to scarcity of physiologically relevant animal models. Indeed, while CMD has been studied in animals with diabetes and/or fed high-fat diets, in fact, hypertension is the predominant risk factor for CMD in humans but is not captured by these models. Thus, we characterized the CMD that arose in female spontaneously hypertensive rats (SHRs). We measured coronary flow reserve, alongside basic cardiovascular function in SHRs and normotensive Wistar Kyoto rats (WKYs) fed a purified diet, as well as SHRs consuming a grain-based chow (GBC) diet. SHRs on a purified diet, but not WKYs or SHRs consuming GBC, exhibited impaired coronary flow reserve as assessed by echocardiography. Thus, SHRs develop a diet-dependent CMD, which can serve as a model to study hypertension-related CMD.

**Graphical Abstract:** 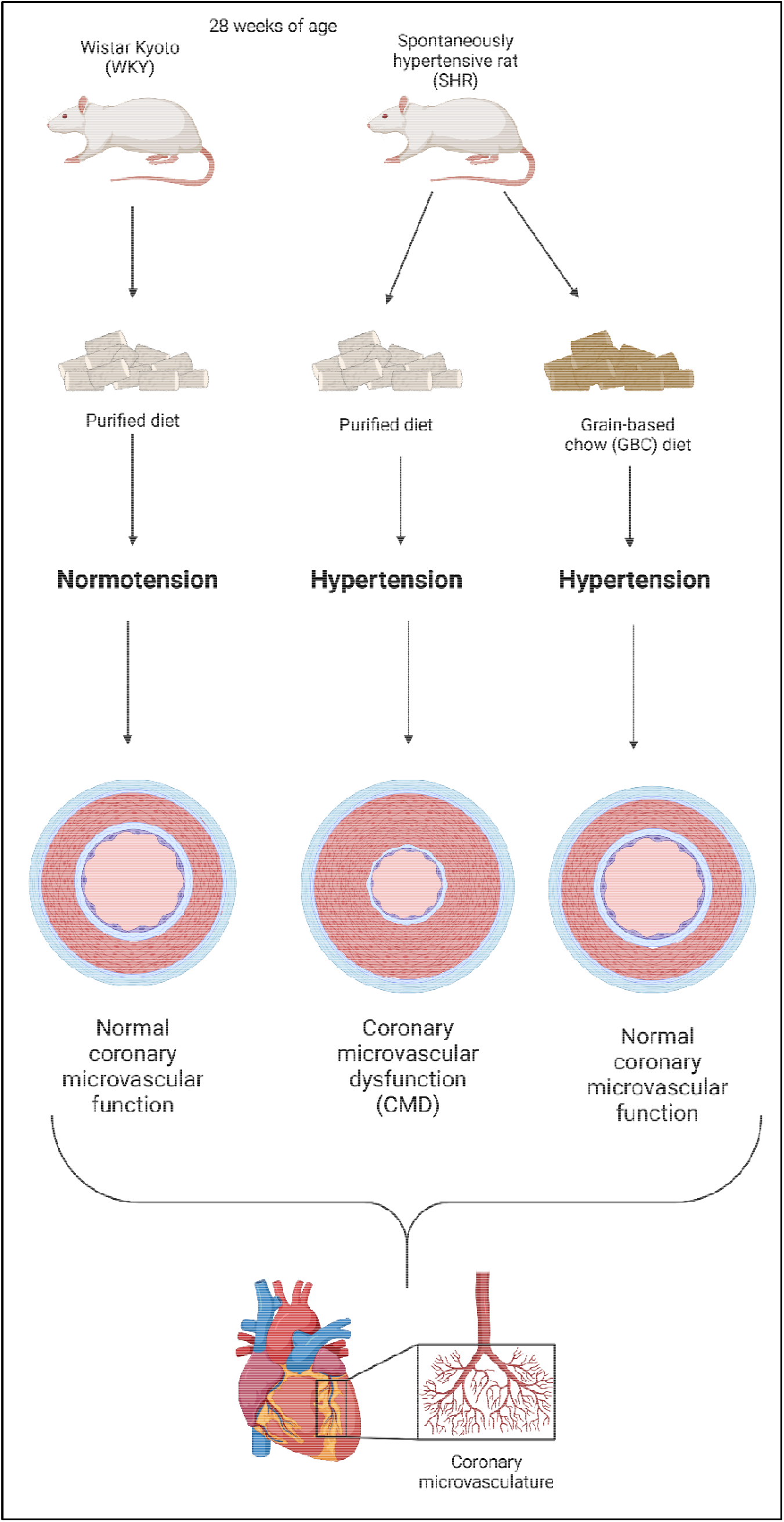

## 1. Introduction

Chest pain (angina pectoris) affects approximately 112 million people globally and is classically associated with ischemic heart disease caused by obstructive atherosclerosis [1]. However, up to two-thirds of women and one-third of men with angina who undergo coronary angiography are found to have no significant epicardial coronary artery obstruction. [2]. This condition is referred to as ischemia with non-obstructive coronary artery disease (INOCA) [3]. INOCA is associated with increased risk of major adverse cardiovascular events, including myocardial infarction and heart failure [3]. Women are also more likely to have INOCA and angina [4], and are four times more likely than men to be re-hospitalized within 6 months following discharge [3]. The underlying pathological driver of INOCA is coronary microvascular dysfunction (CMD), characterized by impaired function of cardiac microvessels which are responsible for approximately 80% of total vascular resistance [5]. CMD is diagnosed by reduced coronary flow reserve (CFR), defined as the ratio of coronary blood flow at maximal hyperemia to that at rest.

Many animal models of CMD have been characterized, but most induce CMD via high-fat diet, hyperlipidemia, or glucose intolerance [6]. While hyperlipidemia and diabetes are indeed associated with CMD, hypertension is a major risk factor for CMD and its prevalence is associated with a stepwise reduction in CFR, while hyperlipidemia is not [7]. In the iPOWER (ImProve diagnOsis and treatment of Women with angina pEctoris and micRovessel disease) study, individuals with the lowest CFR were predominately hypertensive (59%) compared to those with type 2 diabetes (17%) [7]. Thus, currently established animal models do not replicate a clinically prevalent phenotype of CMD driven by hypertension. Spontaneously hypertensive rats (SHRs) develop overt hypertension (SBP>140mmHg) at 6 weeks of age [8], experience progressive cardiac microvascular remodeling as they age [9, 10], and this coincides with significant reductions in CFR [11-13]. Despite these observations, CMD in the SHR has not been traditionally used to study CMD due to limited characterization. Investigators typically utilize male animals which is less clinically relevant in the context of CMD, and dipyridamole is commonly used rather than adenosine, but adenosine is considered a superior, more reliable vasodilator to induce maximal hyperemia for CMD assessment [14]. Further, the impact of diet on microvascular function is typically not considered, despite its potential confounding effects. Thus, the objective of this study was to establish a more clinically relevant hypertensive model of CMD that incorporates sex, diet, and optimized methods for CFR measurement.

## 2. Methods

All animal use and procedures were approved by Georgia State University’s Institutional Animal Care and Use Committee (protocol #: A23025). Three-week-old, female, normotensive Wistar-Kyoto (WKY) rats and SHRs were purchased from Inotiv (West Lafayette, IN, USA). Upon arrival, rats were doubly housed in an environmentally controlled animal care facility and maintained on 12-h light/dark cycles. Rats had free access to water and were maintained on a compositionally-defined open source diet, herein referred to as a “purified diet” (Supplementary table 1) with both soluble (inulin) and insoluble (cellulose) fiber (D23061303; Research Diets). Separately, a group of SHRs consumed standard grain-based chow (GBC; Cat #: 5001, LabDiet, St. Louis, MO). Animals consumed these diets until they were 28 weeks of age.

### 2.1 Blood pressure monitoring

Blood pressure was monitored every two weeks until 24 weeks of age via tail plethysmography using CODA high throughput non-invasive blood pressure system (Kent Scientific, Torrington, CT) in up to four rats simultaneously [15]. All rats were encouraged to walk into the restraint which was adjusted to prevent excessive movement throughout BP recording. The occlusion cuffs were placed at the base of the tail and the volume pressure recording (VPR) cuffs were placed approximately 2 mm adjacent to the occlusion cuffs. Rats rested on the pre-heated heating platform for the duration of each experiment and tail temperatures remained between 35–37 °C. An additional heating blanket was used on top of the restraints for the first 10 minutes of acclimation to warm the animals sufficiently. BP experimental settings were as follows: occlusion cuffs were inflated to 250 mmHg followed by slow deflation over 15 s. The minimum volume changes, as sensed by the VPR cuff, was set to 20 μL. Each recording session consisted of 25 inflation and deflation cycles. During readings, the minimum accepted tail volume was 30 μL. Rats were habituated to the BP measurements over three timepoints before experimental recordings were taken.

### 2.2 Echocardiography

Vevo® 3100 Imaging Platform (Fujifilm Visual Sonics; Toronto, Canada) was used to measure functional and morphological parameters in M-mode with 4-wall LV measurements. Animals were maintained on 2-3% isoflurane on a heated platform. The platform was longitudinally forward angled 5° so that the head was slightly elevated compared to the rest of the body. The ultrasound probe was positioned with a leftward lateral tilt of 60° to obtain the parasternal short axis view. The level of anesthesia was adjusted to obtain a target heart rate of 325 ± 25 beats per minute (bpm). In the prone angle, ophthalmic ointment was used on the eyes to prevent drying. A tourniquet was made using locking forceps and surgical porous tape at the base of the tail to dilate the lateral tail vein. A 26-gauge ¾ inch catheter (Cat #: 07-836-8494, Patterson Veterinary, Loveland, CO) was inserted into the lateral tail vein, the catheter was flushed with sterile PBS, and the catheter was capped to prevent bleeding. The animal was then flipped to the supine position, and hair clippers were used to shave the fur from the neckline to mid chest level. Residual body hair was removed with hair removal cream. Ultrasound transmission gel was applied to the contact area on the rat, and the LV was visualized, first in B-mode. M-mode was used once the largest part of the LV was visualized.

Following LV image capture, the probe was moved cephalically until the aorta, right atrium, and right ventricular outflow tract were visualized (Figure 1A-B), at which point color doppler was utilized to identify coronary flow (Figure 1C). The mode was then changed to pulse wave, and the cursor was placed in the middle of the coronary flow, with minor adjustments to locate the point of maximal flow. The doppler angle was made parallel to the direction of flow, and this was not altered following baseline image capture. Coronary flow velocity is easily identifiable by the “sailboat” pattern of the velocity peaks (Figure 1D). Following baseline coronary velocity capture, the catheter cap was removed, and a sterile 20-gauge 40-inch small bore extension (Cat #: 07-890-7617, Patterson Veterinary, Loveland, CO) was inserted into the catheter. This extension line was pre-filled with a sterile PBS solution containing adenosine at a concentration of 0.5 mg/mL which was attached to a filled 5 mL syringe on an infusion pump (Fusion 100-X Touch Pump, SAI Infusion Technologies, Lake Villa, IL). Infusion proceeded at a rate of 0.25 mg/kg BW per min for 6 min. Coronary flow velocity was monitored during this time, and maximal flow was captured. CFR was calculated as maximal velocity ÷ baseline velocity.

**Figure 1.**
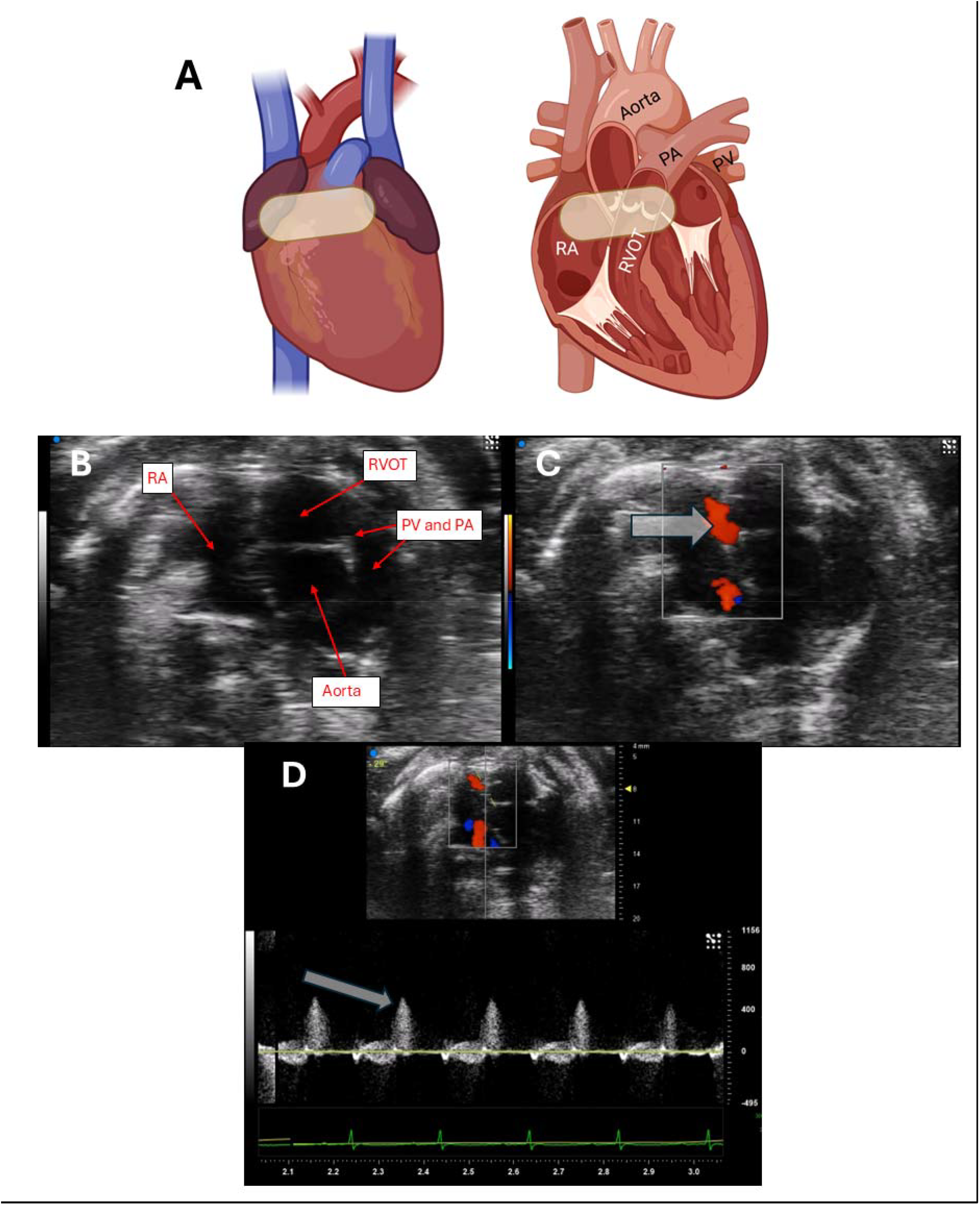
Identification of coronary flow via echocardiography. (A) Location of the echocardiographic probe on the heart, and (B) identification of the structural features. (C) Assessment of coronary flow with color doppler, with the arrow pointing to coronary flow. (D) Pulse wave velocity measurements to quantify coronary flow, with the arrow pointing towards the peak velocity. This velocity pattern resembles a “sailboat”. Abbreviations: PA, pulmonary artery; PV, pulmonary valve; RA, right atrium; RVOT, right ventricular outflow tract. See text for more information.

### 2.3 Nitric oxide metabolite measurements

Whole blood was collected during decapitation. Whole blood was allowed to clot for 40 minutes, placed on ice, then centrifuged at 500 x g for 10 min. Serum was collected from the upper layer, and stored at -80. Nitrates and nitrites were measured fluorometrically with a commercially available kit (Cat #: 780051, Cayman Chemical, Ann Arbor, MI).

### 2.4 Statistical analysis

GraphPad Prism (San Diego, CA) was used for all statistical analyses. Animal data were analyzed using one-way ANOVA followed by Tukey-Kramer post-hoc multiple comparison analysis. Values are represented as mean ± standard deviation (SD). Data were deemed significant if P ≤ 0.05.

## 3. Results and Discussion

Blood pressure was significantly increased in SHRs compared to WKY irrespective of diet (Figure 2). No significant differences were observed between SHRs consuming the purified diet or GBC.

**Figure 2.**
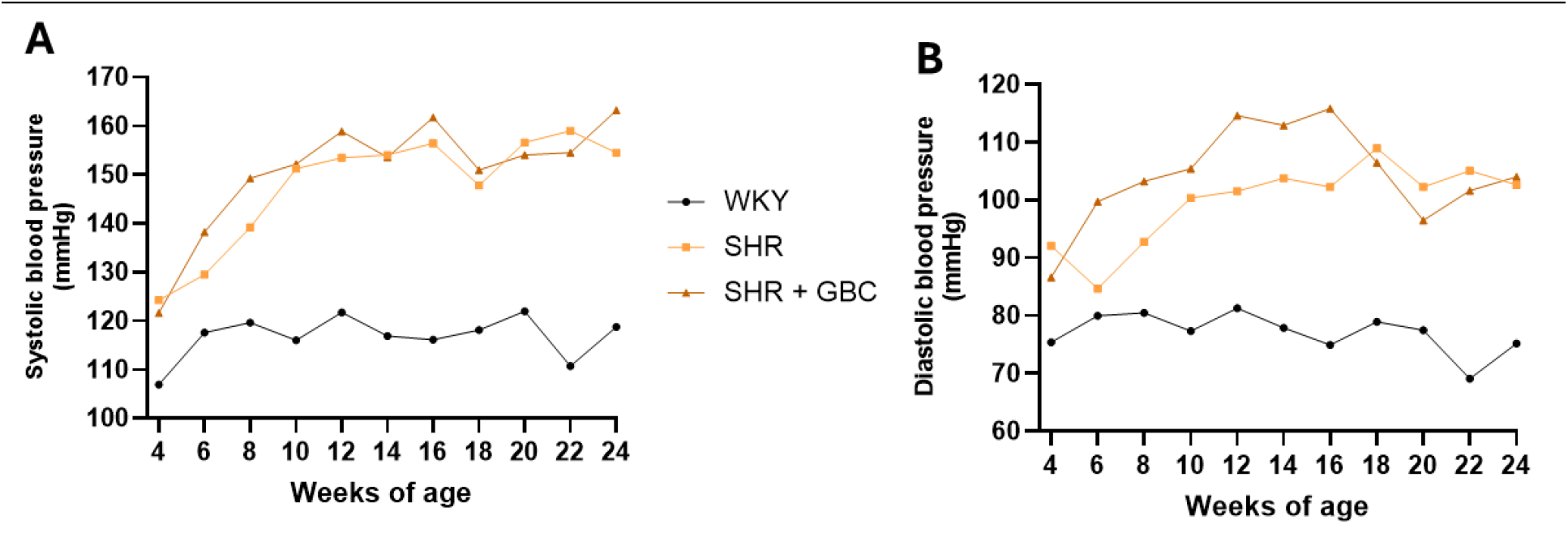
Blood pressure readings during (A) systole and (B) diastole.

SHRs also exhibited cardiac abnormalities irrespective of diet. Specifically, SHRs had lower ejection fraction and fractional shortening, albeit still within the accepted normal range (Fig. 3A-C). This observation is consistent with the literature, and pathologic reductions in EF may occur between 12-18 months of age in males [16] and 24 months in females [17]. Left ventricular mass was significantly reduced in SHRs irrespective of diet, but not when normalized to body weight (Fig. D & E). Thus, LV wall thickness is likely a function of body size. However, the left ventricular internal diameter during systole and diastole was significantly greater in SHRs versus WKY (Fig. 3F & G), suggesting LV dilation.

**Figure 3.**
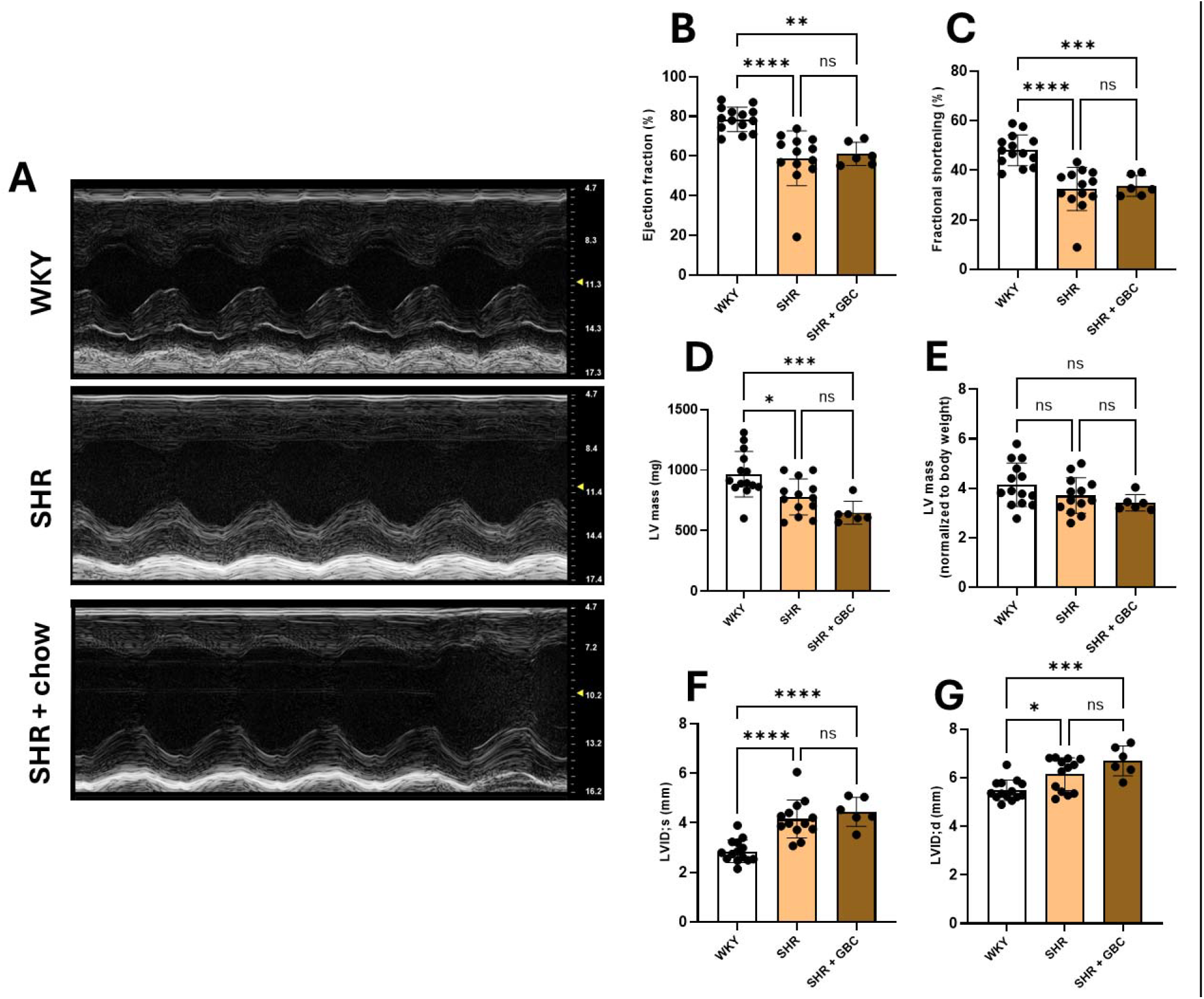
M-mode echocardiographic LV 4-wall measurements. (A) Representative M-mode images of the left ventricle. Quantification of functional parameters, (B) ejection fraction and (C) fractional shortening, as well as morphological parameters (D) LV mass normalized to body weight and (E) LV internal diameter during systole. \**P*≤.05, \*\**P*≤.01, \*\*\**P*≤.001, **** *P*≤.0001.

**Figure 4.**
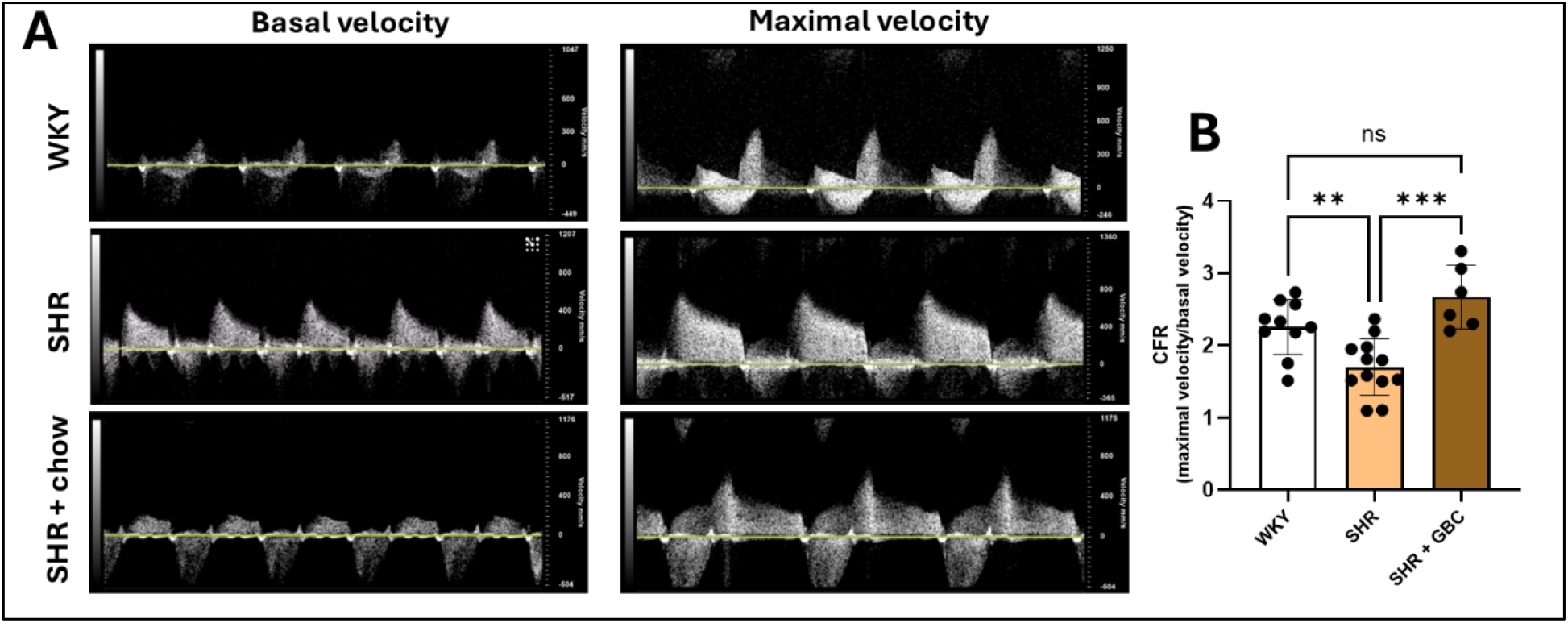
Coronary microvascular function via CFR. (A) Representative pulse wave velocity images prior to and during adenosine infusion (0.25 mg/kg BW per min) and (B) the quantification of CFR. \**P*≤.05, \*\**P*≤.01, \*\*\**P*≤.001, **** *P*≤.0001.

Assessment of cardiac microvascular function revealed that CFR was significantly lower in SHRs compared to age-matched WKY (1.70 ± 0.39 vs 2.25 ± 0.38; *P* = 0.009), both of which had been maintained on the purified diet. In contrast, SHRs consuming GBC displayed CFR levels similar to that of WKYs and significantly greater than that of SHRs fed the purified diet (*P* = 0.0002). Thus, SHRs fed purified diet had impaired coronary microvascular function, while feeding them GBC diet prevented these effects.

Primary ingredients in GBC include corn, soybean, beetroot, fish meal, oats and alfalfa. The purified diet is absent from any plant constituents and can be considered a processed and refined diet. This diet, while low in fat, is more reflective of the human Western diet than GBC, which has traditionally been used in biomedical research. These plant constituents of GBC contain various phytochemicals, including polyphenols and nitrates, which can favorably impact vascular tone [18]. Interestingly, assessment of serum nitrates and nitrites among the three groups did not reveal significant differences (Figure S1), suggesting potentially beneficial effects of the polyphenols from GBC. While these phytochemicals in GBC can reduce blood pressure [19, 20], we did not observe significant differences in blood pressure between SHRs consuming the purified diet and GBC. Thus, there may be beneficial effects in the microvasculature from GBC that are independent of the known deleterious effects of hypertension. Figure 5 highlights the theoretical effects that phytochemicals from diet can have in mediating CMD based on mechanisms we have described previously [21, 22].

**Figure 5.**
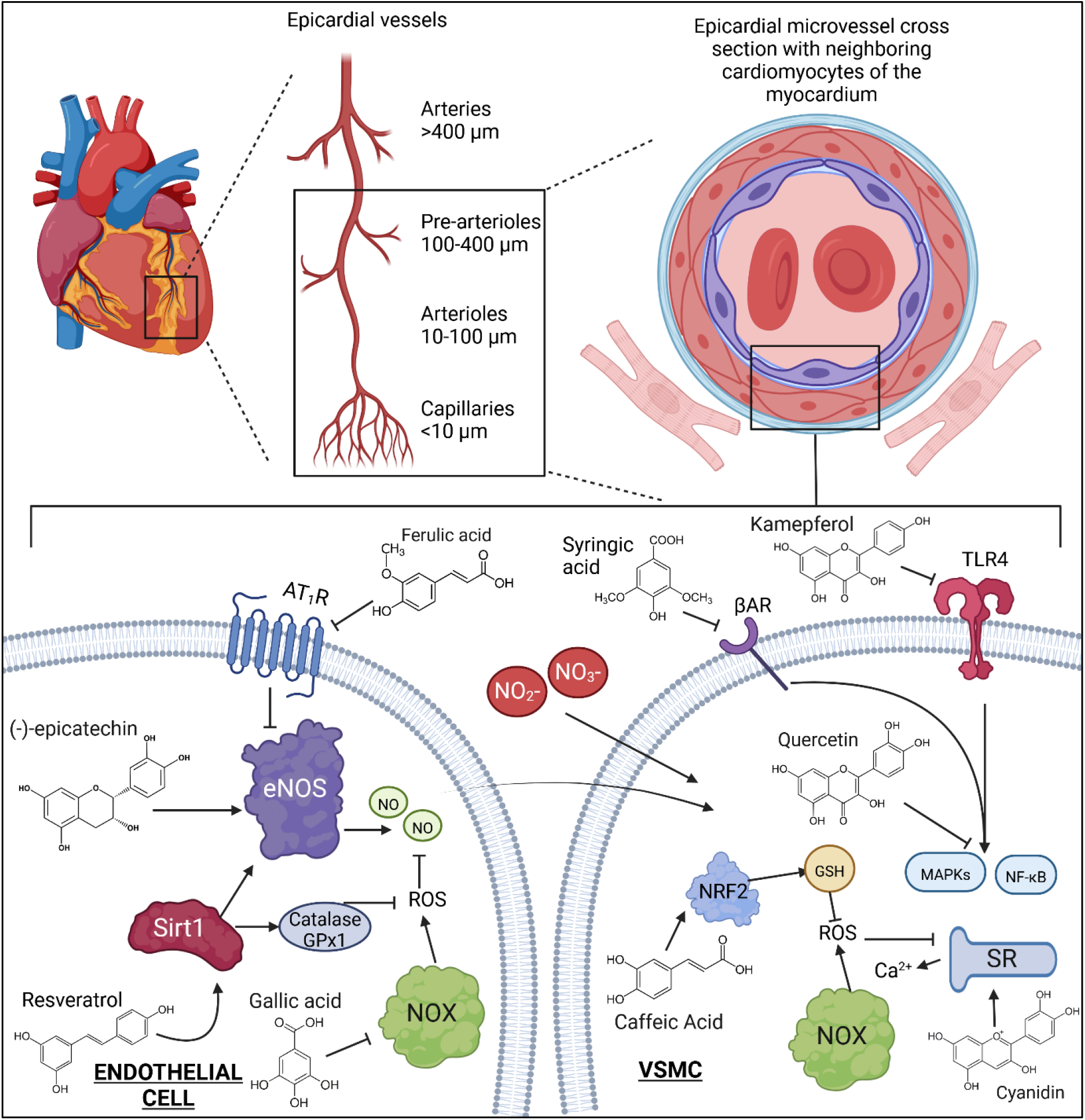
Theoretical molecular mechanisms by which phytochemicals from diet and their metabolites can improve endothelial- and VSMC-dependent cardiac microvascula function [21, 22]. Abbreviations: Angiotensin II type-1 receptor, AT_1_R; Beta adrenergic receptor, βAR; endothelial nitric oxide synthase, eNOS; Glutathione peroxidase-1, GPx1; Glutathione, GSH; Mitogen activated protein kinases, MAPKs; Nuclear factor kappa B, NF-κB; Nuclear factor erythroid 2–related factor 2, NRF2; Nitric oxide, NO; NADPH-oxidase, NOX; Reactive oxygen species, ROS; Sarcoplasmic reticulum, RS; Sirtuin 1, Sirt1; Vascular smooth muscle cell, VSMC.

## 4. Conclusion

In conclusion, a refined, phytochemical-depleted diet induces CMD in female SHRs, but not in normotensive female WKY rats. This hypertensive model of CMD is more reflective of human CMD for which hypertension is a primary risk factor. This model can be used to investigate treatment strategies for CMD or to understand molecular mechanisms of the disease.

## Funding

This work was supported by the Agriculture and Food Research Initiative grant no. 2023-67012-39756/project accession no. 1030574 from the USDA National Institute of Food and Agriculture as well as the National Institutes of Health, grant no. 5R01DK083890-13 and 1R01HL157311.

## Conflicts of Interest

The authors declare no conflicts of interest.

## Notes

### Competing Interest Statement

The authors have declared no competing interest.

