## Supplementary table 1 for "An animal model of coronary microvascular dysfunction (CMD) in the female spontaneously hypertensive rat: the role of diet"

| Table S1. Modified open standard diet composition (D23061303; Research Diets) | |
| --- | --- |
| **Ingredient** | **grams** |
| Casein | 200 |
| L-Cystine | 3 |
| Corn Starch | 252.5 |
| Maltodextrin 10 | 150 |
| Dextrose | 150 |
| Sucrose | 102.41 |
| Cellulose | 75 |
| Inulin | 25 |
| Soybean oil | 70 |
| Mineral Mix S10026A (RD-96 w/o NaCl) | 5 |
| DiCalcium Phosphate | 15.4 |
| Calcium Carbonate | 11 |
| Potassium Citrate, 1 H_2_O | 29 |
| Sodium Chloride | 4.07 |
| Copper Carbonate | 0.0071 |
| Ferric Citrate | 0.074 |
| Zinc Carbonate | 0.022 |
| Vitamin Mix V15937 | 10 |
| Choline Bitartrate | 2 |

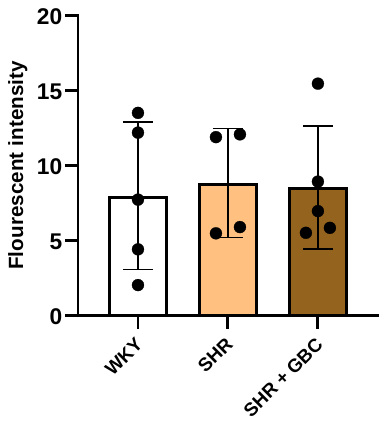

**Figure S1**. Serum nitric oxide metabolite measurements.
